# Maternal transmission as a microbial symbiont sieve, and the absence of lactation in male mammals

**DOI:** 10.1101/2022.01.10.475639

**Authors:** Brennen T. Fagan, George W. A. Constable, Richard Law

## Abstract

Gut microbiomes of humans carry a complex symbiotic assemblage of microorganisms. As in all mammals, the special mode of feeding newborn infants through milk from the mammary gland enhances the opportunity for vertical transmission of the milk microbiome from parents to the gut microbiome of offspring. This has potential benefits, but it also brings with it some hazards for the host. Here we use mathematical and numerical models to demonstrate that vertical transmission from both parents would allow host populations to be invaded by microbiome elements that are deleterious. In contrast, vertical transmission, when restricted to one parent, acts as a sieve preventing the spread of such elements. We show that deleterious symbionts generate selection for uniparental transmission in host populations, and that this selective advantage is maintained in the presence of moderate horizontal transmission. Some vertical transmission from mother to infant is bound to happen in placental mammals. This paper therefore puts forward the hypothesis that the asymmetry between females and males, together with the hazards that come with biparental transmission of the milk microbiome, generate selection against male lactation in humans, and in mammals in general.

---

The absence of male lactation in mammals is a puzzle—there appears to be no universally convincing reason why it should not happen. John Maynard Smith pointed out that paternal care which incorporates such feeding would be evolutionarily stable in monogamous mammals [1]. There have been over 200 My for male lactation to evolve [2]. Genetic control of the mammary gland is widely distributed across mammalian chromosomes [3]. Genetically male mammals, including humans, have the potential to lactate [4–6]. This requires a sufficently high level of the hormone prolactin, which is normally down-regulated in males, preventing lactation from happening [7]. It seems there are selection pressures which keep male lactation firmly switched off.

A well-known answer to the puzzle, building on the work of Trivers [8, 9], is that a bias against paternal care is to be expected when there is uncertainty as to who the father is [10]. The drawback to this explanation is that there are many socially monogamous mammals [1, 11, 12], and high rates of genetic monogamy amongst these species [13]. Since other forms of paternal care have evolved in such species [14], why has male lactation not followed suit? Daly has suggested female lactation might not limit reproductive success [15], but this is questionable because the reproductive cycle would restart sooner if the period of female lactation was shorter [16]. The levels of prolactin needed for lactation do impair male activity [16], and include loss of fertility in human males [17, 18]. However, this is reversible on the restoration of normal prolactin levels [19]. Why should a temporary reduction in male fertility not be evolutionarily sustainable in monogamous mammals? The suppression of lactation in male mammals in general, and male humans in particular remains an open question [16].

We suggest here that vertical transmission of elements of the gut microbiome, near the time at which mammals are born, provides a possible explanation. There is increasing evidence for some vertical transmission of such elements [20, 21], including transmission through breast milk [22]. However, there is a basic, general problem about biparental transmission of symbionts, seen in its strongest form when a rare symbiont first colonises a host population (i.e. when most matings by carriers of the symbiont are with uninfected hosts). When transmission is biparental, it suffices for *either* parent to carry a symbiont for it to be passed on to the offspring. Conversely, when transmission is uniparental, most commonly meaning maternal inheritance, the symbiont is passed on through only one of the parents. The number of hosts with the symbiont in the next host generation under biparental transmission is then double its value under uniparental transmission (Fig. 1). The problem for the host population is that biparental transmission gives the symbionts a reproductive boost, enabling them to invade, even if they are harmful. Uniparental transmission provides an elegant natural solution to this problem, removing the boost and leaving the frequency of vertically transmitted symbionts unchanged when hosts reproduce. This solution has been recognised in other areas of the literature since early work on the evolution of uniparental cytoplasmic inheritance [23, 24], and its extension to the more general context of symbiosis [25].

**FIG. 1.**
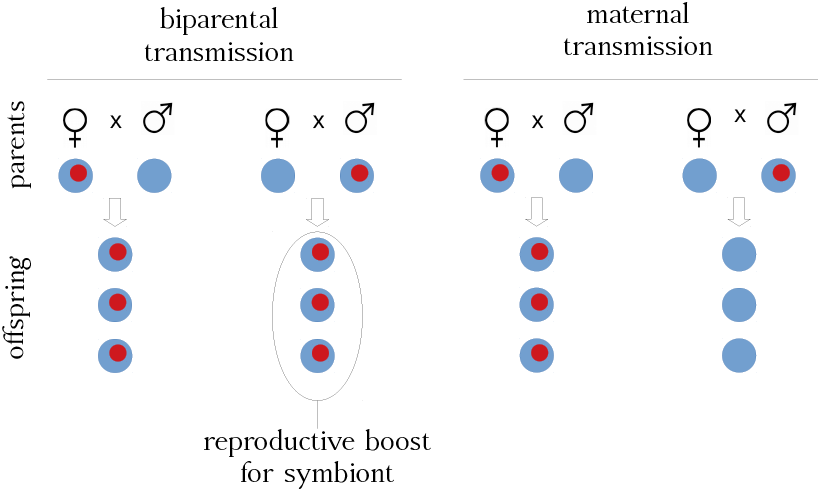
Biparental transmission in a host population gives a reproductive boost to symbionts. This is at its greatest when symbionts (red) are rare in the host population, and infected hosts mate mostly with uninfected ones, as shown here. The boost is prevented by uniparental transmission, assumed here to be maternal.

## THE SYMBIONT SIEVE

The advantage to the host of restricting symbiont transmission to one rather than both parents can be made precise with a little algebra, previously used for investigating the evolution of anisogamy [24]. Consider a host’s symbiont community comprising a set of microbial taxa, labelled *s*. Suppose an additional element appears and changes the set of symbionts (e.g. a new microbial taxon is added to the original community, *s*), such that this new set is labelled *S*. We examine how the fate of *S* is determined by the mode of vertical transmission, and by the effect of *S* on host fitness. In doing this we make two important simplifying assumptions: that the new symbiont is independent of resident microbiome, and that there is no evolution in the microbiome. Interactions in the microbiome are of course important [26] and so is its evolution [27, 28], but are not needed to illustrate the simple basic effects of uniparental transmission on the microbiome, and the feedback from the microbiome to evolution of vertical transmission that we describe here. The frequency of matings under random mating, and the resulting symbiont communities associated with these matings, are given in Table I. In this scheme, the frequency of hosts carrying symbiont community *S* (respectively *s*) in the parental generation is *p* (respectively *q* = 1 − *p*). For generality, we write the probability that a female (respectively male) passes *S* on to the next generation as *α* (respectively *β*). Thus, under biparental transmission *α* = 1, *β* = 1, and under maternal transmission *α* = 1, *β* = 0. *P*(*S*) is the probability with which the offspring of the mating hosts carry the symbiont community *S*, and *P*(*s*) is the probability with which the offspring carry the symbiont community *s*.

**TABLE I.** Matings with two host types, one type with symbiont community *s* and the other with community *S* augmented by a new symbiont, and the associated probabilities of the offspring’s community, as explained in the text.

| mother | × | father | frequency | $P(S)$ | $P(s)$ |
| --- | --- | --- | --- | --- | --- |
| $s$ | × | $s$ | $q^2$ | 0 | 1 |
| $S$ | × | $s$ | $pq$ | $\alpha$ | $1 - \alpha$ |
| $s$ | × | $S$ | $pq$ | $\beta$ | $1 - \beta$ |
| $S$ | × | $S$ | $p^2$ | $\alpha + \beta - \alpha\beta$ | $(1 - \alpha)(1 - \beta)$ |

The frequency of hosts carrying *S* in the next generation, *p*^*′*^, is given by the frequency of matings that pass *S* on, multiplied by the fitness *w* of hosts carrying *S* relative to those those carrying *s*:

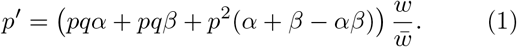

Here the mean fitness of the population, 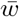, normalises the frequencies in the next generation so that they sum to 1. The change in frequency of hosts carrying *S* is then

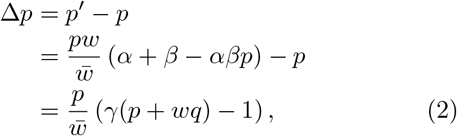

where *γ* = *α* + *β αβp*.

The condition for the symbiont community *S* to increase in the host population from one generation to the next is Δ*p >* 0. From (2), the condition is (*p*+*wq*) *>* 1*/γ*. When *S* first appears in the host population, *p* ≈ 0 and *q* ≈ 1, so *S* invades if *w >* 1*/γ*. Fully maternal transmission of the symbionts (*α* = 1, *β* = 0) implies *γ* = 1, so *S* can only invade if it gives a host fitness *w >* 1, i.e. a fitness greater than that of *s*. However, fully biparental transmission of symbionts (*α* = 1, *β* = 1) implies *γ* = 2, so *S* invades if it gives a host fitness *w >* 1*/*2 relative to *s*, i.e. a fitness potentially lower than that of *s*. This demonstrates the reproductive boost that allows symbionts to invade under biparental transmission, even if they reduce host fitness. Finally, with *w <* 1*/*2, *S* would not invade at all in this simple model, whether transmission is uniparental or biparental.

A similar argument could be constructed to show that transmission of symbionts through the father (*α* = 0, *β* = 1) rather than the mother, would be just as advantageous. However, the process of birth makes some transmission of microorganisms from the mother to infant unavoidable in placental mammals. Evidence for this includes differences observed in the gut microbiomes of infants born naturally, and those delivered by caesarean section [29], which implies some transfer of symbionts occurs during a natural birth [30]. There are also indications that bacteria can reach the uterus through the blood stream of the mother in mice [31], although the longstanding paradigm of the sterile womb in humans still has some support [32, 33]. Given this basic asymmetry between the roles of mother and father in placental mammals, uniparental transmission of symbionts would need to operate through the mother.

In effect, maternal transmission acts as a first line of defense for hosts in the face of an unruly mob of microorganisms. It operates at the start of life as a “symbiont sieve”, separating those that are beneficial to the host (e.g. mutualistic microbes) from those that are deleterious (e.g. pathogenic microbes), and preventing the deleterious ones from spreading in the host population.

Fig. 2 illustrates the sieve in action. This shows the ultimate fate of symbiont communities augmented by a new element (community *S*) giving host fitness *w* relative to hosts without it (*s*), under biparental and maternal transmission. To do this, we constructed a stochastic, birth-death model to describe host population dynamics, controlling for vertical transmission of the symbiont community (see). The effect of the new symbiont on host fitness *w* was taken to be a random variable drawn from a (truncated) normal distribution centred on *w* = 1, the point of neutrality where there is no effect on host fitness (Fig. 2a). For argument’s sake, the symbiont acted on the host death rate *d* as a factor *d/w*, being beneficial to the host when *w >* 1, and deleterious when *w <* 1. We carried out 5000 independent trials of this experiment for each mode of vertical transmission. Each trial started with a host population close to its equilibrium population size (1000 hosts), 10 of which carried the new symbiont (community *S*), and the population was tracked over time to see whether the symbiont was ultimately present in all or none of the hosts.

**FIG. 2.**
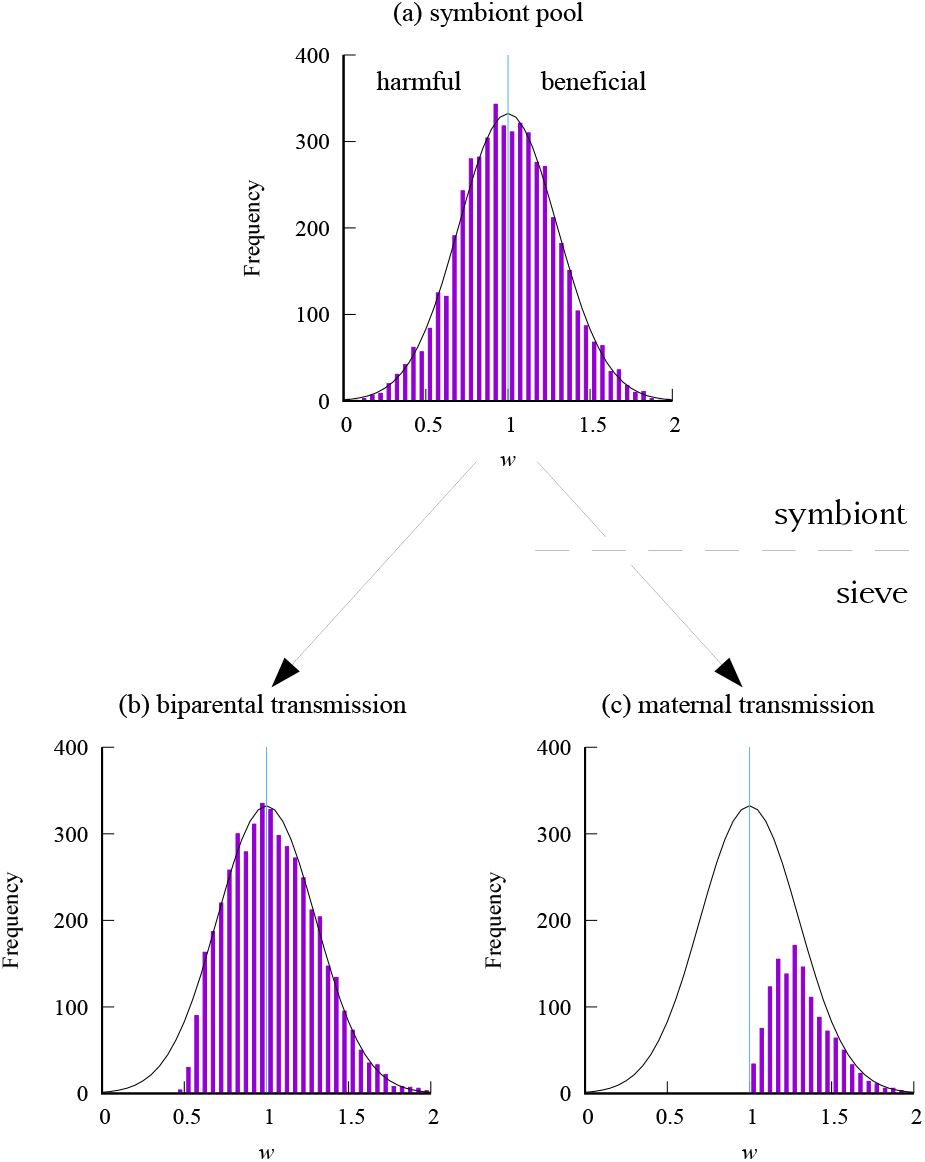
A numerical experiment to demonstrate maternal transmission operating as a symbiont sieve. (a) A frequency distribution from which to draw random values of *w*, the effect of symbiont community *S* on host fitness relative to community *s*; *w <* 1: *S* harmful; *w >* 1: *S* beneficial. (b) Frequency distribution of *w* in symbionts that successfully invade under biparental transmission: invasion can occur whether the symbiont is beneficial or harmful to the host. (c) Corresponding frequency distribution under maternal transmission: this sieves out harmful symbionts, only allowing invasion by symbionts beneficial to their hosts.

Fig. 2b shows that the strong boost from biparental transmission allows invasion by deleterious symbionts, as long as they no more than double the death rate of hosts. In other words, the lower limit for invasion by the new symbiont is *w* = 1*/*2, as in the simpler algebraic model above. In contrast, maternal transmission (Fig. 2c) operates as a sieve, preventing invasion by deleterious symbionts (*w <* 1), while still allowing the beneficial ones to invade (*w >* 1), as in the algebraic model above. Near the point of neutrality under maternal transmission (*w* = 1), demographic stochasticity is likely to lead to extinction of the symbiont before it can get established. In an infinitely large population, the probability of invasion would tend to zero as *w →* 1 from above.

Fig. 3 gives some examples of time series on which Fig. 2 is based, taking specific values of *w* to show how the outcomes differ under biparental and maternal transmission. The threshold for invasion by community *S* carrying the new symbiont is *w*_0_ = 1*/*2 for biparental and *w*_0_ = 1 for maternal transmission (see). Thus Fig. 3a, b with *w* = 0.2 show the symbiont being rapidly eliminated from the host population, irrespective of the mode of transmission. However, the outcomes are quite different at *w* = 0.6 because the symbiont can invade under biparental but not under maternal transmission (Fig. 3c, d). At *w* = 1, where the symbiont has no effect on host fitness, the reproductive boost from biparental transmission still allows rapid invasion by the symbiont, whereas its fate is the outcome of an unbiased random walk under maternal transmission (Fig. 3e, f). It is only above both invasion thresholds that the symbiont can eventually spread to all hosts (i.e. symbiont frequency = 1) under both modes of transmission (Fig. 3g, h).

**FIG. 3.**
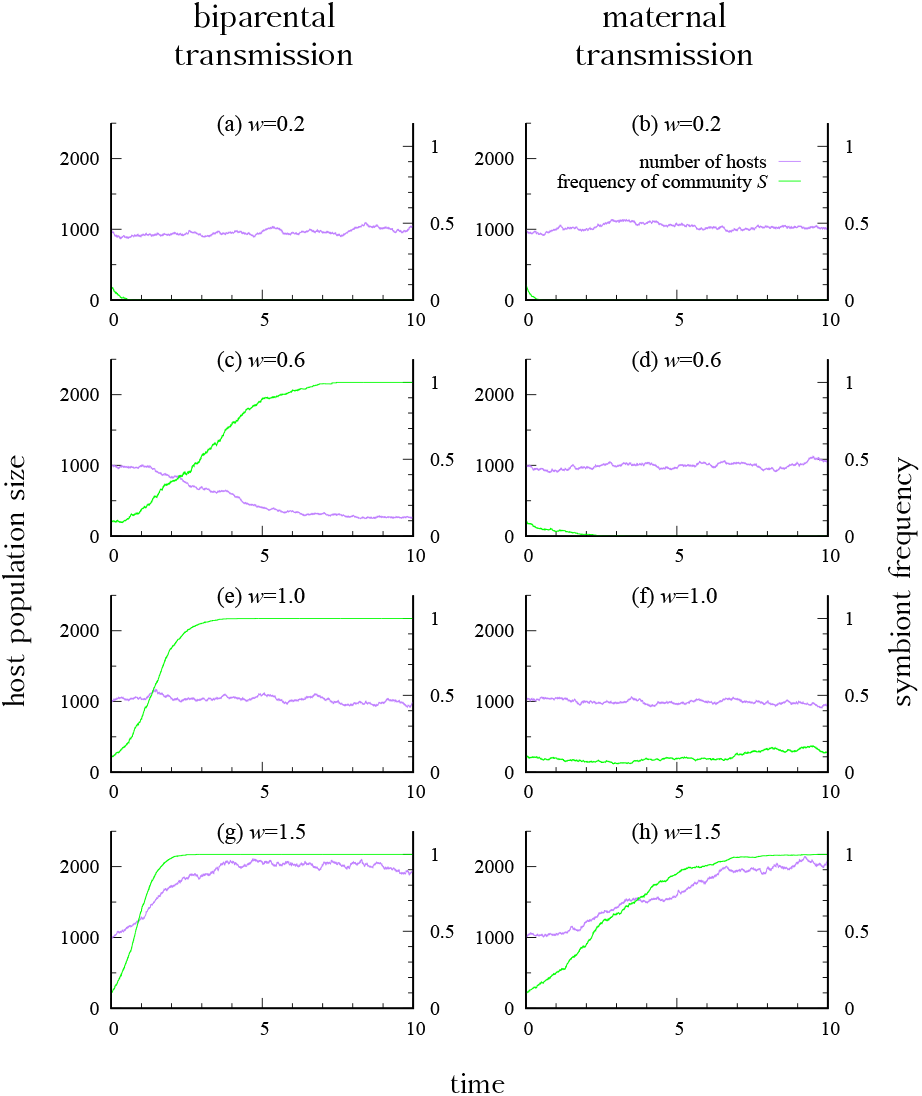
Example time-series of host population size, starting with 1000 individuals of which 100 had community *S* carrying the new symbiont (i.e. a starting frequency of 0.1 for ease-of-visualisation). In column 1 transmission of the symbiont is biparental; in column 2 it is maternal. Rows show different values of *w* (the effect on host fitness), from the top where symbionts are harmful, to the bottom where they are beneficial. Maternal transmission protects host populations from invasion by deleterious symbionts which can occur under biparental transmission.

Notice though that the protection afforded by maternal transmission comes with the cost that the symbiont spreads more slowly than it would under biparental transmission (Fig. 3g, h). How significant an issue this is depends on the distribution of host fitnesses generated by the pool of available symbionts. The distribution is unknown, but it most likely skewed towards symbionts with deleterious effects (*w <* 1), after a sequence of successful invasions has taken place. By this stage, protection from deleterious symbionts is likely to be most important. The successional sequence through which the microbiome develops is a matter for future research, e.g. using the methods of adaptive dynamics [34, 35].

## EVOLUTION OF THE SYMBIONT SIEVE

In the previous section, we have seen that a population of hosts transmitting symbionts maternally is more effective at suppressing emergent deleterious symbionts than a host population transmitting symbionts biparentally. However, how host genetic systems evolve and maintain maternal transmission of symbionts is a separate and interesting matter. As a deleterious symbiont is spreading under biparental transmission (Fig. 3c), it is present in some hosts and absent in others. If neither mother nor father carry the symbiont, host fitness is unaffected by the mode of transmission. The same applies if both mother and father already carry the symbiont. However, if the father carries the symbiont and the mother does not, a gene that stops transmission from the father gains an advantage, as we show in Fig. 4.

**FIG. 4.**
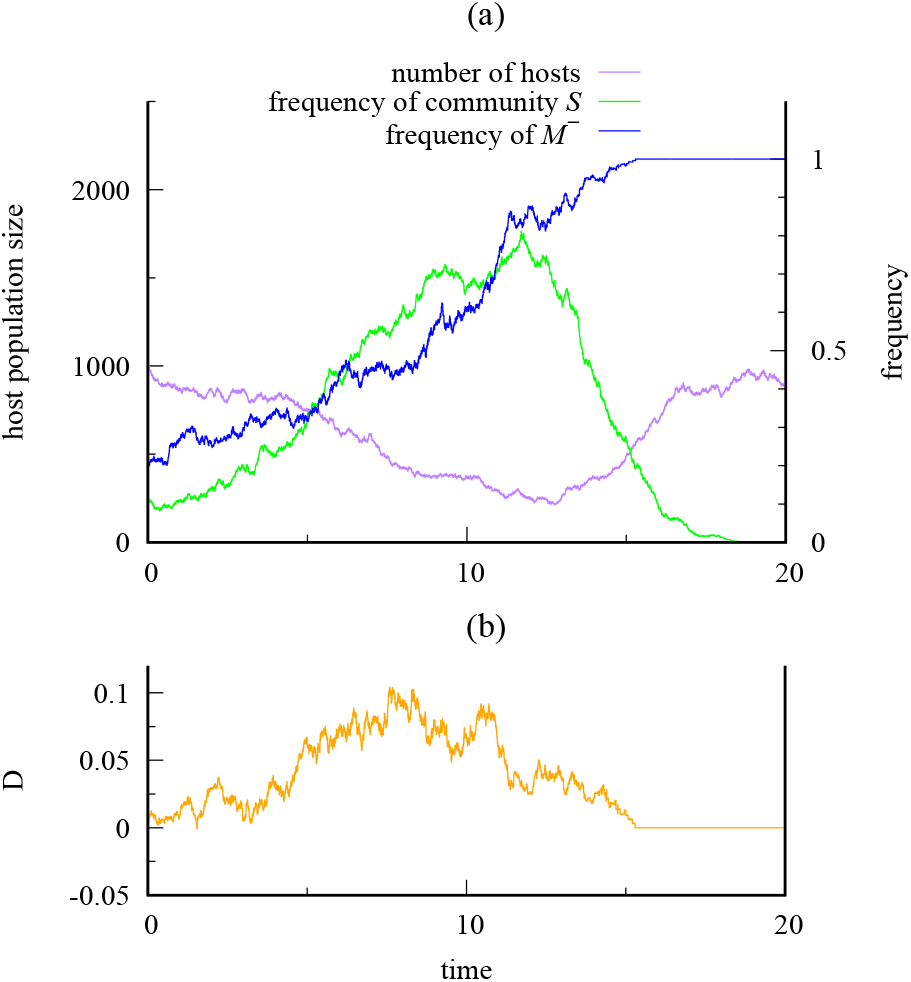
A harmful symbiont (*w* = 0.6) drives evolution of maternal transmission. (a) Here a host population starts mostly with biparental transmission (high frequency of host gene *M* ^+^, low frequency of *M* ^−^), and a low frequency of the symbiont community *S* in the population. This allows the new symbiont to spread. The symbiont generates a selective advantage for the *M* ^−^ gene which prevents male transmission. The host population then becomes dominated by maternal transmission *M* ^−^, and under these conditions the symbiont cannot persist. (b) The gene *M* ^−^ gains its advantage by becoming associated with male hosts that lack the symbiont, as measured by the coefficient of disequilibrium *D*.

To illustrate evolution of maternal transmission, Fig. 4 uses the numerical model in the, with two alternative genes {*M* ^+^, *M* ^−^*}* to control vertical transmission by males in the host population, during an invasion by a deleterious symbiont (*w* = 0.6). (Transmission through females is always present.) For simplicity, we assumed the genes were at a locus on the Y chromosome. *M* ^+^ allows transmission from males, making vertical transmission biparental, whereas *M* ^−^ prevents transmission from males, making vertical transmission maternal.

Starting with a low frequency of *M* ^−^, a deleterious symbiont can invade the host population, because vertical transmission is predominantly biparental (Fig. 4a). In turn, *M* ^−^ increases in frequency, because it becomes associated with hosts lacking the symbiont which are fitter than those carrying the symbiont. This association can be measured by a coefficient of disequilibrium *D* (see), which becomes positive (Fig. 4b). The association drives the host population towards maternal transmission, and eventually *M* ^−^ reaches a frequency great enough to turn the tables against the deleterious symbiont. *M* ^−^ goes to fixation, making vertical transmission fully maternal, and the deleterious symbiont is then eliminated from the host population. The outcome is a host population with both maternal transmission and protection from harmful symbionts with 1 *> w >* 1*/*2.

An alternative scenario (not shown here) is that *M* ^−^ does not reach fixation before the symbiont is present in all hosts. In this case, the path to maternal transmission goes in steps: first *M* ^−^ increases whenever a community containing a deleterious symbiont is present; then on fixation (or loss) of the deleterious symbiont, *M* ^−^ and *M* ^+^ are neutral with respect to one another (there is only one community present in the population, so the mode of transmission carries no fitness effects); selection for *M* ^−^ is triggered again by the emergence of a new deleterious symbiont that generates a new community *S*^*′*^ and so on. In this way maternal transmission is fixed in the host population under recurrent introductions of symbionts that are relatively deleterious with respect to the resident.

Evolution from biparental to maternal transmission sheds some new light on host-symbiont evolution. It is not vertical transmission per se that checks the spread of deleterious symbionts in host populations with separate sexes. Rather it is the restriction of vertical transmission to just one sex, usually females, that makes the symbiont sieve work. This restriction emerges from natural selection driven by deleterious symbionts themselves, and the pattern is so ubquitous in nature [30] that it is usually taken as given in research on the theory of symbiont transmission [27, 36–40].

## HORIZONTAL TRANSMISSION AND THE SYMBIONT SIEVE

It needs to be kept in mind that hosts are still vulnerable to invasion by deleterious symbionts if horizontal transmission, which allows hosts to pick up symbionts from the environment, is also present. We examine this in Fig. 5, extending our model to allow a fixed per-capita rate *e*_0_ *>* 0 at which hosts acquire the new symbiont from an environmental source [40] (note we do not account for direct horizontal transmission between hosts). When *e*_0_ is sufficiently large, horizontal transmission floods any controls generated by vertical transmission. But when *e*_0_ is lower (of the same order as birth and death rates), vertical transmission still exerts some control over the symbiont community, and the outcome then depends on whether vertical transmission is biparental or maternal (uniparental).

**FIG. 5.**
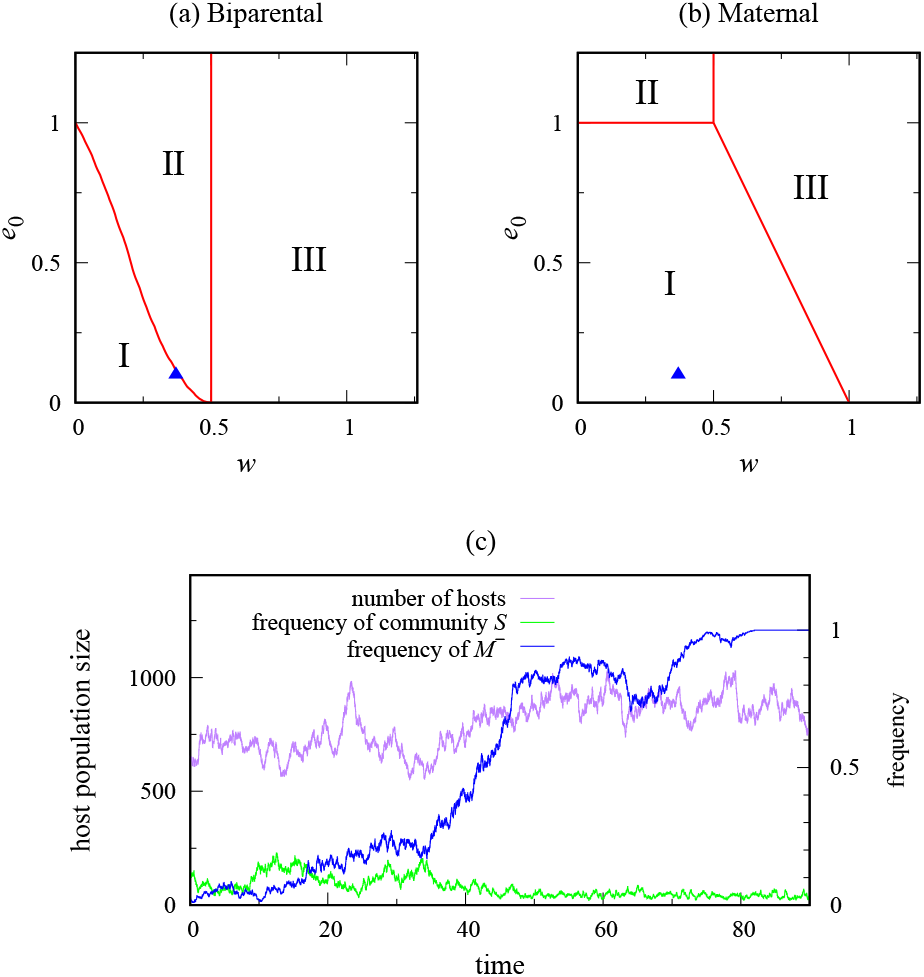
Three contrasting outcomes (I, II, III) following arrival of a new symbiont when horizontal transmission occurs, in addition to (a) biparental and (b) maternal vertical transmission. Horizontal transmission takes place at a fixed percapita rate *e*0, and *w* is the effect of the symbiont on host fitness. Region I: the host population remains bimorphic: some hosts carry the symbiont, and others do not. Region II: host population goes to extinction. Region III: the new symbiont is present in all hosts. (c) Bimorphism allows selection for a gene *M* ^−^ causing loss of male transmission; this shifts host vertical transmission from biparental to maternal. The reverse path from maternal to biparental transmission is not possible for *w <* 1. The example uses *w* = 0.37, *e*0 = 0.1, shown as a triangle in (a) and (b). Outcomes in (a) and (b) were computed by numerically solving a system of ordinary differential equations (see) and categorising the parameters by the number and type of fixed points produced. The example path in (c) was computed by simulation.

Specifically in Fig. 5, Region I of the space (*w, e*_0_) prevents a deleterious symbiont from going to fixation. Instead, the host population goes to a bimorphic equilibrium at which there are two host types present: those that carry the new harmful symbiont, and those that do not. This is important because host genes modifying vertical transmission need both types of host to be present for natural selection to act on them. Horizontal and biparental transmission together (Fig. 5a, Region I) then result in selection for the modifier gene *M* ^−^ that switches off transmission through males, and the host population evolves to a state of horizontal and maternal transmission (Fig. 5b), as illustrated by the path in Fig. 5c. There is no return route to biparental transmission in Region I because the modifier gene *M* ^−^ benefits from the bimorphic state. Horizontal transmission in effect helps to protect maternal transmission.

Note that there is a Region II in Fig. 5a in which the reproductive boost to the symbiont from biparental transmission, coupled with its harmful effect on the host, drives the host population to extinction. This region does not exist under maternal transmission for *e*_0_ *<* 1, and emphasizes the extra dangers resulting from biparental transmission in the presence of deleterious symbionts.

In Region III in Figs. 5a and 5b, the new symbiont does go to fixation, and the subset *w <* 1 of this region is the range over which the successful symbiont is harmful to the host. This subset is independent of *e*_0_ under biparental transmission, and includes *e*_0_ = 0. However, Region III gets smaller as *e*_0_ decreases under maternal transmission, and the symbiont sieve fully protects the host population when transmission is strictly vertical (i.e. *e*_0_ = 0). Horizontal transmission dominates when *e*_0_ ≥ 1, irrespective of whether vertical transmission is biparental or maternal. Also, beneficial symbionts (*w >* 1) go to fixation irrespective of the type of vertical transmission, and they do so faster when this transmission is biparental.

To summarise, we recall that Region I in Fig 5a is the parameter regime in which maternal transmission will be selected for when first emerging in a population with biparental transmission of the microbiome. Having reached fixation, maternal transmission will resist re-invasion by biparental transmission in the parameter regime denoted by Region I in Fig 5b. Although Region I may seem restrictive as plotted, we note that this is a reasonable place for an observed evolved system to find itself in. The alternatives are the elimination of the host species (Region II), or overwhelming environmental and selective pressure that eliminates diversity between host microbiomes in the population (Region III).

## DISCUSSION

There are many ways in which hosts can limit vertical transmission of microorganisms from the father, through physiology, behaviour and social structure. Human breast milk nicely illustrates one of them. Breast milk contains a large microbiome [41, 42]. This becomes a major feature in the composition of the infant’s gut microbiome [29], and there is little doubt that some vertical transmission of the microbiome is taking place. The source of the microbiome is a subject of current research. But at least some elements appear to come from the maternal gut, possibly reaching the breast through an entero-mammary pathway [43], and are transmitted from mother to infant via breast milk [44]. Deleterious elements would spread through host populations if both parents took part in breast feeding. In practice, the hormone prolactin is down-regulated in males to a level low enough to stop lactation. This has the immediate effect of greatly restricting the contribution fathers can make to the gut microbiome of newborn infants.

We cannot prove that the danger of invasion by deleterious elements in the gut microbiome is the reason why there is no male lactation. So this argument is put forward as a hypothesis. However, milk production for feeding infants is a defining property of mammals and lactation by males is almost invariably switched off in them. The symbiont sieve thus offers a novel solution to this longstanding puzzle, and points to an important mechanism by which mammals exert some control over the composition of their gut microbiomes during transmission of microorganisms from one generation to the next. The hypothesis offers a new perspective on several lines of research.

First, it is testable. Widespread deleterious symbionts propagating solely through vertical transmission in breast milk of humans and other mammals would be inconsistent with the maternal symbiont sieve. This is not to claim that there are no maternally transmitted pathogens, which is clearly incorrect, but such symbionts would need additional mechanisms to spread, such as direct horizontal infection from host to host. Indeed, although in humans there are numerous examples of viruses that can be transmitted through breast milk (including human immunodeficiency virus and cytomegalovirus [45]), there does not appear to be as yet any evidence of deleterious viruses for which breast milk is the primary transmission mode.

Secondly, it leads to the question as to what beneficial roles are played by maternally transmitted symbionts in gut microbiomes. These are open systems colonized within host generations by a diverse set of microorganisms, and are of remarkable complexity [26]. The order in which these systems are assembled leaves a lasting impact on the structure of the microbiome in mice [46], suggesting a key role for those present near the time of birth. For example, *Bifidobacterium*, which occurs in breast milk, is known to inhibit the growth of pathogenic bacteria and to aid in the digestion of the milk [47], with strong evidence for the vertical transfer and control of these bacteria during breast feeding [48, 49].

Thirdly, mammalian gut microbiomes are notable for the way in which their structure mirrors the phylogenetic relationships of their mammalian hosts (phylosymbiosis), notwithstanding their species richness. This is thought to be related to special traits of mammals including viviparous birth, milk production and parental care [21]. The role of physiological filters by hosts on their symbionts is recognised in this, but not the basic property of maternal transmission in separating beneficial from harmful symbionts. It could be that positive frequency-dependent selection between beneficial symbionts and their hosts acts to conserve microbiome structure, contributing to the signal of host phylogeny.

Fourthly are the exceptions. The Dayak fruit bat is one of the very few mammals in which males with functional mammary glands have been documented under natural conditions [4]. Gut microbiomes of bats are unlike those of other mammals, being quite unpredictable (as in birds), and lacking clear relationships with host phylogeny [50]. This could be in part a consequence of adaptations for flight, including lower body mass, a reduction gut size and shorter retention times of materials in the gut. It is possible that the selection pressures associated with flight have led to a different evolutionary path in the Dayak fruit bat, resulting in both parents to contributing to feeding [1].

Developing a theory of vertical transmission requires a recognition that both biparental and uniparental transmission could occur, and that they have strikingly different consequences for microbiota associated with the hosts. Future work will need to deal with the role the symbiont sieve plays in the makeup of the gut microbiome (not dealt with here). This calls for an understanding of the ecology of the microbial communities, moving on from neutral models [37] and those with frequency-independent selection [27, 51], to take account of priority effects, succession and development of invasion resistance [52–54], which have measurable effects on host fitness [55]. Generalised Lotka-Volterra models [56, 57] and consumer-resource models [58, 59] are a step in this direction. In addition, the feedback from the microbiome to host fitness can evidently drive host evolution, selecting host genes that control vertical transmission of symbionts. The key modelling challenge here is how to map properties of the microbiome (e.g. presence or absence of particular species and their abundances) to emergent fitness effects on hosts, which may even depend on host genotype [28, 60].

The evolutionary question we have tackled in this paper is quite different from a major current focus of host-microbiome theory, namely whether a symbiont will evolve to increase host fitness at the expense of its own fitness within the microbiome [27]. Our question is how the gut microbiome operates as a driver of evolution of vertical transmission in mammalian hosts. Although this is just one of many points of contact between hosts and their microbiomes, it is especially important in setting the path along which the microbiome develops in the host offspring [61], with effects that carry over to their subsequent health [29]. Our focus on evolution of vertical transmission also offers a new perspective on horizontal transmission, over which hosts have relatively little direct control. Horizontal transmission is thought to decrease the potential for coevolution between hosts and their microbiomes [27]. However, our results show the presence of deleterious symbionts in the host population (sustained through horizontal transmission) actively holds in place the symbiont sieve through a continuing selection pressure against host modifier genes that would allow transmission from both parents.

Finally, we note that maternal transmission of symbionts is a widespread phenomenon in the natural world [30]. We suggest that maternal transmission could have evolved a number of times as a simple mechanism that has the effect of sieving beneficial from harmful symbionts during vertical transmission.

## METHODS

To investigate the dynamics of a host population with vertically transmitted symbionts, we constructed a continuous-time, stochastic, birth-death process with logistic density-dependence. This is more nuanced than the algebraic model in the text; the density dependence allows a wider range of outcomes, including extinction of the host driven by deleterious symbionts. The state of the host population at time *t* consisted of the sex {♀, ♂} and the presence or absence {+, −} of an added symbiont in each host individual *i* = 1, …, *n*(*t*), i.e.:

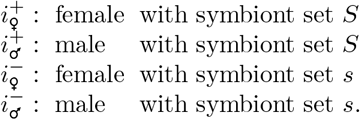

We write *n*^+^(*t*) as the number of hosts carrying the new symbiont (community *S*), and *n*^−^(*t*) as the number without this symbiont (community *s*). Thus *n*^+^(*t*) gives a measure of the abundance of the additional symbiont in the host population at time *t*, and *n*^+^(*t*)*/n*(*t*) is its frequency in the host population.

The probability per unit time of death *d*_*i*_ of host *i* contained a logistic component common to all individuals, and a further dependence on symbiont status:

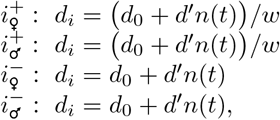

where *d*_0_ is an intrinsic death rate and *d*^*′*^ scales the density-dependent component of death. The death rate was modified by a factor 1*/w* in hosts carrying the additional symbiont (symbiont community *S*). This describes the effect of the new symbiont on its host: *w* = 1 is neutral, a beneficial symbiont (*w >* 1) lowers *d*_*i*_ by a factor 1*/w*, and a deleterious symbiont (0 *< w <* 1) raises *d*_*i*_ by a factor 1*/w*.

The per-female probability per unit time of giving birth *b*_0_ was set to be independent of host population density. The sex of the newborn individual was assigned with an equal probability 0.5 to be female or male. Thus the probability per unit time with which a mother gave birth to a daughter, the key measure for host population growth, was *b*_0_*/*2. Whether the symbiont was present (+) or absent (–) in a newborn host individual depended on the mode of vertical transmission:

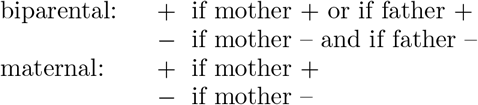

Mating was assumed to be at random, so a random father was chosen from the males present in the population for biparental transmission. This completes the specification of the stochastic, birth-death process.

We carried out realisations of the stochastic process using the Gillespie algorithm [62]. Simulated results were obtained with parameter values: *b*_0_ = 4, *d*_0_ = 1, *d*^*′*^ = 0.001. The computations for Fig. 2 used a random value of *w* drawn from a normal distribution with mean 1, and standard deviation 0.3 (truncated at 0 and 2.5). The initial number of host individuals was 1000 (the equilibrium point in the absence of the new symbiont), and the new symbiont was introduced to 10 individuals at the start, randomly distributed between females and males. Realisations were terminated when the symbionts were present in all hosts, or absent in all hosts, or the run-time had reached 200 time units. 5000 independent realisations of each mode of vertical transmission were carried out. In almost all instances the outcome was presence in all hosts, or absence in all hosts. The four exceptions out of 10000 realisations were under maternal transmission, with *w* close to neutrality.

We extended the stochastic process above to describe evolution of vertical transmission using two alternative genes at a locus on the Y chromosome. Male hosts were classified according to the gene they carried, *M* ^+^ switching on male transmission, and *M* ^−^ switching off male transmission. Symbiont transmission from females was present throughout, so vertical transmission was biparental in crosses with *M* ^+^ males and maternal in crosses with *M* ^−^ males. There are four classes of males depending on transmission gene *{M* ^+^, *M* ^−^*}* and presence or absence of the new symbiont *{*+, −*}*. To measure the association between transmission gene and symbiont status, we write the frequency of the classes in males as

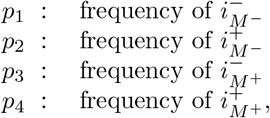

where *p*_1_ + *p*_2_ + *p*_3_ + *p*_4_ = 1. The coefficient of disequilibrium, *D* = *p*_1_*p*_4_ − *p*_2_*p*_3_, then measures the association between transmission gene and symbiont status. *D* is positive if the new symbiont is under-represented in *M* ^−^ males and over-represented in *M* ^+^ males, and negative if vice versa. This completes the specification of the stochastic, birth-death process with evolution of vertical transmission. Computation of Fig. 4 was carried out with parameter values set to be the same as those in Figs 2 and 3.

We note that a system of ordinary differential equations can be constructed for the mean behaviour of the stochastic process. We have used this to check the results of the stochastic realisations, to gain further understanding of the dynamics, and to check the robustness of the results to changes in parameter values. In the absence of evolution (Figs 2, 3), the associated equations are:

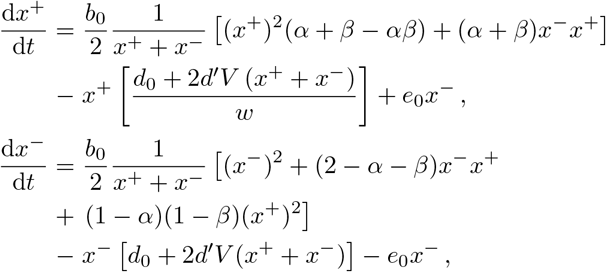

assuming a 1:1 sex ratio to reduce the dimensionality of the system from four to two. The state variables are the density of female hosts *x*^+^ with the symbiont community *S*, and the density *x*^−^ with the community *s*. These state variables come from dividing the number of females with and without the additional symbiont (*n*^+^, *n*^−^) by a scaling parameter *V* (*x*^+^ = *n*^+^*/V, x*^−^ = *n*^−^*/V*). The mating classes are as defined in Table I, the proportion of females (respectively males) passing community *S* on to the next host generation being *α* (respectively *β*). We included an extra term to allow horizontal transmission, using a fixed per-capita rate *e*_0_ at which hosts encounter the additional symbiont in the environment and switch the symbiont community from *s* to *S*. Under strict vertical transmission, *e*_0_ = 0.

We used the differential equations to describe the dynamics under biparental transmission (setting *α* = 1, *β* = 1), and under maternal transmission (setting *α* = 1, *β* = 0). The additional symbiont was added close to a boundary equilibrium point *I* of the host population

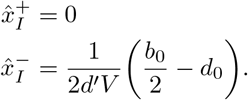

The initial per-capita rate of increase of hosts carrying the symbiont at this equilibrium point is

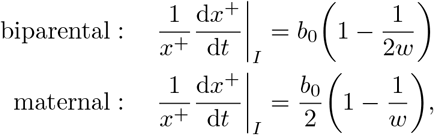

giving a threshold fitness *w*_0_ above which invasion of the symbiont happens with values *w*_0_ = 1*/*2 for biparental transmission, and *w*_0_ = 1 for maternal transmission. A second equilibrium point *II* occurs at

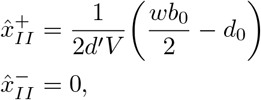

where every host carries the symbiont. This is unaffected by the mode of transmission, but, unless the symbiont is neutral (*w* = 1), the symbiont changes the equilibrium host population size when it is present in all hosts. There is a threshold fitness *w*_1_ at which the symbiont causes extinction of the host population 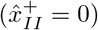 at *w*_1_ = 2*d*_0_*/b*_0_.

When horizontal transmission is absent, the thresholds for invasion by the symbiont *w*_0_ and extinction of the host population *w*_1_ are *w*_0_ = *w*_1_ = 0.5 under biparental transmission, and *w*_0_ = 1, *w*_1_ = 0.5 under maternal transmission, with the parameter values used here. Thus maternal transmission protects the host population from extinction, but biparental transmission could bring the host population size close to zero.

Numerical analysis was conducted using the above system of differential equations in Mathematica. We varied the parameters *w* and *e*_0_ between 0 and 1.5 in steps ranging from 0.001 to 0.1 and, at each combination of parameter values, the system was numerically solved and each physical (*x*^+^, *x*^−^ ≥ 0) solution’s stability was calculated. This procedure was used to calculate the dividing lines between the regions in Figs. 5a and 5b, which were then verified using simulations. We then repeated this procedure for our robustness check for values of *b*_0_ between 2 and 6 in steps of 0.25 and *d*_0_ between 0.5 and 2 in steps of 0.125. This robustness check revealed that the majority of cases are as described in the main text with some differences as to the locations and curvature of the boundary lines. There is also the possibility of overlap between Regions I and III in which the final state of the population depends on the initial conditions. As our focus in the main text is on invasions of either the new symbiont or maternal transmission, this would result in domination of the overlap by Region I (due to the small *x*^+^ initial condition).

## FUNDING AND ACKNOWLEDGEMENTS

B.T.F. is funded by a Leverhulme Trust Research Centre the Leverhulme Centre for Anthropocene Biodiversity. We thank the University of York Reading Group on Stability and Complexity in which discussions on the dynamics of gut microbiomes led to this work, J. A. Fagan for useful discussions, and C. M. Law and J. W. Pitchford for comments on the paper.

## Code Archival

Mathematica and C code are available at https://github.com/Brennen-Fagan/Maternal-Transmission.

## Notes

### Competing Interest Statement

The authors have declared no competing interest.

### Summary of Updates

This version has been amended to extend the evolutionary arguments put forward in the original paper, to account for horizontal transmission, and to address a broader range of the relevant biological literature.

## References

[1] J. Maynard Smith, Parental investment: a prospective analysis, Animal Behaviour 25, 1 (1977).

[2] C. M. Lefevre, J. A. Sharp, and K. R. Nicholas, Evolution of lactation: Ancient origin and extreme adaptations of the lactation system, Annual Review of Genomics and Human Genetics 11, 219–238. doi:10.1146/annurev (2010).

[3] D. Lemay, D. Lynn, and W. e. a. Martin, The bovine lactation genome: insights into the evolution of mammalian milk, Genome Biol. 10, R43 (2009).

[4] C. M. Francis, E. L. P. Anthony, J. A. Brunton, and T. H. Kunz, Lactation in male fruit bats, Nature 367, 691 (1994).

[5] T. Reisman and Z. Goldstein, Case report: Induced lactation in a transgender woman, Transgender Health 3, 24 (2018).

[6] J. García-Acosta, R. San Juan-Valdivia, F.-M. A.D., N. Lorenzo-Rocha, and M. Castro-Peraza, Trans* pregnancy and lactation: A literature review from a nursing perspective., Int J Environ Res Public Health. 17, 44 (2019).

[7] L. O’Hara, M. Curley, M. Tedim Ferreira, L. Cruickshanks, L. Milne, and L. B. Smith, Pituitary androgen receptor signalling regulates prolactin but not gonadotrophins in the male mouse, PLOS ONE 10, 1 (2015).

[8] R. L. Trivers, Parental investment and sexual selection, in Sexual Selection and the Descent of Man, edited by B. Campbell (Aldine Publishing, Chicago, 1972) p. 136–179.

[9] R. L. Trivers, Parental investment and sexual selection (Routledge, 2017).

[10] H. Kokko and M. D. Jennions, Parental investment, sexual selection and sex ratios, Journal of evolutionary biology 21, 919 (2008).

[11] D. G. Kleiman, Monogamy in mammals, The Quarterly review of biology 52, 39 (1977).

[12] D. Lukas and T. H. Clutton-Brock, The evolution of social monogamy in mammals, Science 341, 526 (2013).

[13] T. H. Clutton-Brock and K. Isvaran, Paternity loss in contrasting mammalian societies, Biology Letters 2, 513 (2006).

[14] C. Kvarnemo, Why do some animals mate with one partner rather than many? a review of causes and consequences of monogamy, Biological Reviews 93, 1795 (2018).

[15] M. Daly, Why don’t male mammals lactate?, Journal of theoretical Biology 78, 325 (1979).

[16] T. H. Kunz and D. J. Hosken, Male lactation: why, why not and is it care?, Trends in Ecology & Evolution 24, 80 (2009).

[17] C. Carani, A. R. M. Granata, M. F. Fustini, and P. Marrama, Prolactin and testosterone: their role in male sexual function, International Journal of Andrology 19, 48 (1996), https://onlinelibrary.wiley.com/doi/pdf/10.1111/j.1365-2605.1996.tb00434.x.

[18] W. Hair, O. Gubbay, H. Jabbour, and G. Lincoln, Prolactin receptor expression in human testis and accessory tissues: localization and function, Molecular human reproduction 8, 606 (2002).

[19] C. Ufearo and O. Orisakwe, Restoration of normal sperm characteristics in hypoprolactinemic infertile men treated with metoclopramide and exogenous human prolactin, Clinical Pharmacology & Therapeutics 58, 354 (1995).

[20] A. H. Moeller, T. A. Suzuki, M. Phifer-Rixey, and M. W. Nachman, Transmission modes of the mammalian gut microbiota, Science 362, 453 (2018).

[21] E. K. Mallott and K. R. Amato, Host specificity of the gut microbiome, Nature Reviews Microbiology 19, 639 (2021).

[22] S. Duranti, G. A. Lugli, C. Milani, K. James, L. Mancabelli, F. Turroni, G. Alessandri, M. Mangifesta, W. Mancino, M. C. Ossiprandi, A. Iori, C. Rota, G. Gargano, S. Bernasconi, F. Di Pierro, D. van Sinderen, and M. Ventura, Bifidobacterium bifidum and the infant gut microbiota: an intriguing case of microbe-host co-evolution, Environmental microbiology 21, 3683 (2019).

[23] I. M. Hastings, Population genetic aspects of deleterious cytoplasmic genomes and their effect on the evolution of sexual reproduction, Genetical Research 59, 215 (1992).

[24] R. Law and V. Hutson, Intracellular symbionts and the evolution of uniparental cytoplasmic inheritance, Proceedings of the Royal Society of London Series B 248, 69 (1992).

[25] S. A. Frank, Host–symbiont conflict over the mixing of symbiotic lineages, Proceedings of the Royal Society of London. Series B: Biological Sciences 263, 339 (1996).

[26] K. R. Foster, J. Schluter, K. Z. Coyte, and S. Rakoff-Nahoum, The evolution of the host microbiome as an ecosystem on a leash, Nature 548, 43 (2017).

[27] S. Van Vliet and M. Doebeli, The role of multilevel selection in host microbiome evolution, Proceedings of the National Academy of Sciences 116, 20591 (2019).

[28] J. Roughgarden, S. F. Gilbert, E. Rosenberg, I. Zilber-Rosenberg, and E. A. Lloyd, Holobionts as units of selection and a model of their population dynamics and evolution, Biological Theory 13, 44 (2018).

[29] C. J. Stewart, N. J. Ajami, J. L. O’Brien, D. S. Hutchinson, D. P. Smith, M. C. Wong, M. C. Ross, R. E. Lloyd, H. V. Doddapaneni, G. A. Metcalf, D. Muzny, R. A. Gibbs, T. Vatanen, C. Huttenhower, R. J. Xavier, M. Rewers, W. Hagopian, J. Toppari, A.-G. Ziegler, J.-X. She, B. Akolkar, A. Lernmark, H. Hyoty, K. Vehik, J. P. Krischer, and J. F. Petrosino, Temporal development of the gut microbiome in early childhood from the TEDDY study, Nature 562, 583–588. doi:10.1038/s41586 (2018).

[30] L. J. Funkhouser and S. R. Bordenstein, Mom knows best: the universality of maternal microbial transmission, PLoS Biol 11, e1001631. doi:10.1371/journal.pbio.1001631 (2013).

[31] E. Jiménez, M. L. Mariń, R. Martiń, J. M. Odriozola, M. Olivares, J. Xaus, L. Ferńandez, and J. M. Rodríguez, Is meconium from healthy newborns actually sterile?, Research in Microbiology 159, 187 (2008).

[32] M. E. Perez-Muñoz, M.-C. Arrieta, A. E. Ramer-Tait, and J. Walter, A critical assessment of the “sterile womb” and “in utero colonization” hypotheses: implications for research on the pioneer infant microbiome, Microbiome 5, 48. doi:10.1186/s40168 (2017).

[33] M. C. de Goffau, S. Lager, U. Sovio, F. Francesca Gaccioli, E. Cook, S. J. Peacock, J. Parkhill, D. S. Charnock-Jones, and S. G. C. S, Human placenta has no microbiome but can contain potential pathogens, Nature 572, 329 (2019).

[34] U. Dieckmann and R. Law, The dynamical theory of coevolution: a derivation from stochastic ecological processes, Journal of mathematical biology 34, 579 (1996).

[35] É. Kisdi, A. Stefan, and H. Geritz, Adaptive dynamics: a framework to model evolution in the ecological theatre, Journal of mathematical biology 61, 165 (2010).

[36] M. Lipsitch, S. Siller, and M. Nowak, The evolution of virulence in pathogens with vertical and horizontal transmission, Evolution 50, 1729 (1996).

[37] Q. Zeng, J. Sukumaran, S. Wu, and A. Rodrigo, Neutral models of microbiome evolution, PLoS computational biology 11, e1004365 (2015).

[38] P. T. Leftwich, M. P. Edgington, and T. Chapman, Transmission efficiency drives host–microbe associations, Proceedings of the Royal Society B 287, 20200820 (2020).

[39] J. Roughgarden, Holobiont evolution: Population genetic theory for the hologenome, bioRxiv., doi.org/10.1101/2020.04.10.036350 (2021).

[40] R. Zapién-Campos, F. Bansept, M. Sieber, and A. Traulsen, On the effect of inheritance of microbes in commensal microbiomes, BMC ecology and evolution 22, 1 (2022).

[41] K. M. Hunt, J. A. Foster, L. J. Forney, U. M. E. Schütte, D. L. Beck, Z. Abdo, L. K. Fox, J. E. Williams, M. K. McGuire, and M. A. McGuire, Characterization of the diversity and temporal stability of bacterial communities in human milk, PLoS ONE 6, e21313. doi:10.1371/journal.pone.0021313 (2011).

[42] L. F. Stinson, A. S. M. Sindi, A. S. Cheema, C. T. Lai, B. S. Mühlh äusler, M. E. Wlodek, M. S. Payne, and D. T. Geddes, The human milk microbiome: who, what, when, where, why, and how?, Nutrition Reviews 79, 529–543. doi:10.1093/nutrit/nuaa029 (2020).

[43] G. Oikonomou, M. F. Addis, C. Chassard, M. E. F. Nader-Macias, I. Grant, C. Delbes, C. I. Bogni, Y. Le Loir, and S. Even, Milk microbiota: What are we exactly talking about?, Frontiers in Microbiology 11:60, doi:10.3389/fmicb.2020.00060 (2020).

[44] T. Jost, C. Lacroix, C. P. Braegger, F. Rochat, and C. Chassard, Vertical mother–neonate transfer of maternal gut bacteria via breastfeeding, Environmental Microbiology 16, 2891–2904. doi:10.1111/1462 (2014).

[45] A. E. Palmquist, M. T. Perrin, D. Cassar-Uhl, K. D. Gribble, A. B. Bond, and T. Cassidy, Current trends in research on human milk exchange for infant feeding, Journal of Human Lactation 35, 453 (2019).

[46] I. Martínez, M. X. Maldonado-Gomez, J. C. Gomes-Neto, H. Kittana, H. Ding, R. Schmaltz, P. Joglekar, R. J. Cardona, N. L. Marsteller, S. W. Kembel, et al., Experimental evaluation of the importance of colonization history in early-life gut microbiota assembly, Elife 7, e36521 (2018).

[47] S. Arboleya, C. Stanton, C. A. Ryan, E. Dempsey, and P. R. Ross, Bosom buddies: the symbiotic relationship between infants and bifidobacterium longum ssp. longum and ssp. infantis. genetic and probiotic features, Annual review of food science and technology 7, 1 (2016).

[48] H. Makino, A. Kushiro, E. Ishikawa, D. Muylaert, H. Kubota, T. Sakai, K. Oishi, R. Martin, K. Ben Amor, R. Oozeer, et al., Transmission of intestinal bifidobacterium longum subsp. longum strains from mother to infant, determined by multilocus sequencing typing and amplified fragment length polymorphism, Applied and environmental microbiology 77, 6788 (2011).

[49] S. Duranti, G. A. Lugli, L. Mancabelli, F. Armanini, F. Turroni, K. James, P. Ferretti, V. Gorfer, C. Ferrario, C. Milani, et al., Maternal inheritance of bifidobacterial communities and bifidophages in infants through vertical transmission, Microbiome 5, 1 (2017).

[50] S. J. Song, J. G. Sanders, F. Delsuc, J. Metcalf, K. Amato, M. W. Taylor, F. Mazel, H. L. Lutz, K. Winker, G. R. Graves, G. Humphrey, J. A. Gilbert, S. J. Hackett, K. P. White, H. R. Skeen, S. M. Kurtis, J. Withrow, T. Braile, M. Miller, K. G. McCracken, J. M. Maley, V. O. Ezenwa, A. Williams, J. M. Blanton, V. J. McKenzie, and R. Knight, Comparative analyses of vertebrate gut microbiomes reveal convergence between birds and bats, mBio 11, e02901 (2020).

[51] R. Zapién-Campos, M. Sieber, and A. Traulsen, The effect of microbial selection on the occurrence-abundance patterns of microbiomes, Journal of the Royal Society Interface 19, 20210717 (2022).

[52] K. Shea and P. Chesson, Community ecology theory as a framework for biological invasions, Trends in Ecology & Evolution 17, 170 (2002).

[53] H. M. Kurkjian, M. J. Akbari, and B. Momeni, The impact of interactions on invasion and colonization resistance in microbial communities, PLoS Computational Biology 17, e1008643 (2021).

[54] E. J. Stevens, K. A. Bates, and K. C. King, Host microbiota can facilitate pathogen infection, PLoS pathogens 17, e1009514 (2021).

[55] E. R. Davenport, J. G. Sanders, S. J. Song, K. R. Amato, A. G. Clark, and R. Knight, The human microbiome in evolution, BMC biology 15, 1 (2017).

[56] K. Z. Coyte, J. Schluter, and K. R. Foster, The ecology of the microbiome: networks, competition, and stability, Science 350, 663 (2015).

[57] D. Gonze, K. Z. Coyte, L. Lahti, and K. Faust, Microbial communities as dynamical systems, Current opinion in microbiology 44, 41 (2018).

[58] A. Goyal, T. Wang, V. Dubinkina, and S. Maslov, Ecology-guided prediction of cross-feeding interactions in the human gut microbiome, Nature communications 12, 1 (2021).

[59] P.-Y. Ho, B. H. Good, and K. C. Huang, Competition for fluctuating resources reproduces statistics of species abundance over time across wide-ranging microbiotas, Elife 11, e75168 (2022).

[60] J. Roughgarden, Holobiont evolution: Population theory for the hologenome, bioRxiv 10.1101/2020.04.10.036350 (2022), https://www.biorxiv.org/content/early/2022/05/13/2020.04.10.0363

[61] S. A. Frese, D. A. Mackenzie, D. A. Peterson, R. Schmaltz, T. Fangman, Y. Zhou, C. Zhang, A. K. Benson, L. A. Cody, F. Mulholland, N. Juge, and J. Walter, Molecular characterization of host-specific biofilm formation in a vertebrate gut symbiont, PLoS genetics 9, e1004057 (2013).

[62] D. T. Gillespie, Exact stochastic simulation of coupled chemical reactions, The journal of Physical Chemistry 81, 2340 (1977).

